# Receptor compaction and GTPase movements drive cotranslational protein translocation

**DOI:** 10.1101/2020.01.07.897827

**Authors:** Jae Ho Lee, SangYoon Chung, Yu-Hsien Hwang Fu, Ruilin Qian, Xuemeng Sun, Shimon Weiss, Shu-ou Shan

## Abstract

Signal recognition particle (SRP) is a universally conserved targeting machine that couples the synthesis of ~30% of the proteome to their proper membrane localization^1,2^. In eukaryotic cells, SRP recognizes translating ribosomes bearing hydrophobic signal sequences and, through interaction with SRP receptor (SR), delivers them to the Sec61p translocase on the endoplasmic reticulum (ER) membrane^1,2^. How SRP ensures efficient and productive initiation of protein translocation at the ER is not well understood. Here, single molecule fluorescence spectroscopy demonstrates that cargo-loaded SRP induces a global compaction of SR, driving a >90 Å movement of the SRP•SR GTPase complex from the vicinity of the ribosome exit, where it initially assembles, to the distal site of SRP. These rearrangements bring translating ribosomes near the membrane, expose conserved Sec61p docking sites on the ribosome and weaken SRP’s interaction with the signal sequence on the nascent polypeptide, thus priming the translating ribosome for engaging the translocation machinery. Disruption of these rearrangements severely impairs cotranslational protein translocation and is the cause of failure in an SRP54 mutant linked to severe congenital neutropenia. Our results demonstrate that multiple largescale molecular motions in the SRP•SR complex are required to drive the transition from protein targeting to translocation; these post-targeting rearrangements provide potential new points for biological regulation as well as disease intervention.

---

Mammalian SRP is a ribonucleoprotein complex composed of six proteins (SRP19, SRP9/14, SRP68/72, and SRP54) bound to the 7SL RNA^3^. The universally conserved SRP54 contains an M-domain that binds the 7SL RNA and recognizes the signal sequence or transmembrane domain (TMD) on the nascent polypeptide emerging from the ribosome exit tunnel. A special GTPase domain in SRP54, termed NG, forms a GTP-dependent complex with a homologous NG-domain in SR, and the two NG-domains reciprocally stimulate each other’s GTPase activity^4,5^. SRP is essential^1,6^, and mutations in SRP54-NG cause severe congenital neutropenia with Shwachman-Diamond-like features^7,8^, but the molecular basis of the defect is unknown. Previous works have primarily focused on how SRP recognizes its cargo and assembles with SR^4,5,9,10^. These works showed that signal sequence-bearing ribosome-nascent chain complexes (RNCs) reposition SRP54-NG to dock at uL23 near the ribosome exit site, and SRP pre-organized in this ‘Proximal’ conformation is optimized for assembly with SR^5,9,11^ to deliver translating ribosomes to the ER (Fig. 1a, left and Fig. 1b).

**Figure 1.**
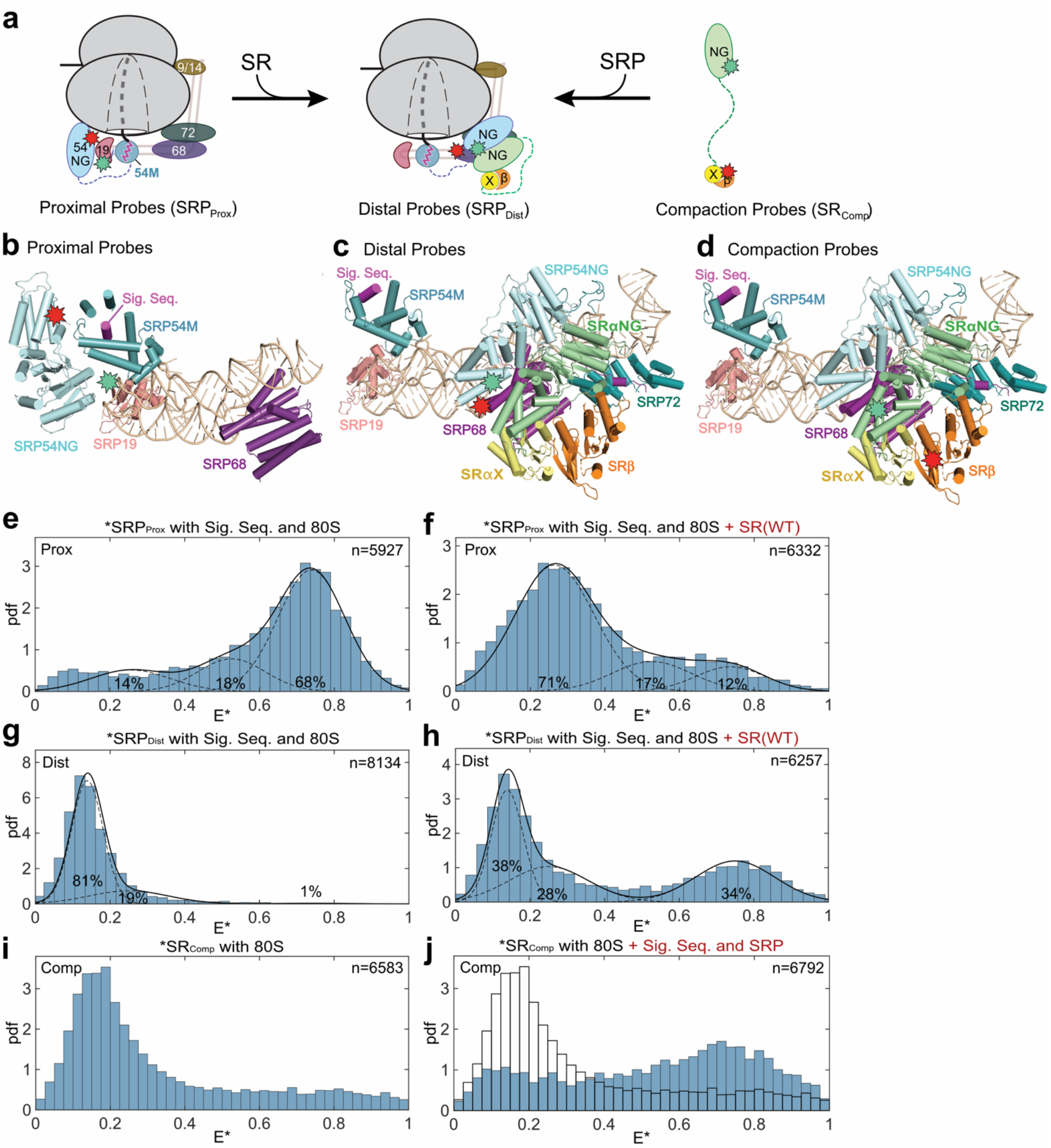
smFRET measurements detect multiple conformational changes upon SRP-SR assembly. **a**, Schematic of conformational changes in SRP and SR indicating the locations of FRET donor (*green*) and acceptor (*red*) dyes. Left, the ‘Proximal’ conformation of SRP in which SRP54-NG is bound near the ribosome exit site. Middle, the ‘Distal’ conformation in which SRP54-NG docks near SRP68/72 at the distal site. Right, the third FRET pair to measure the end-to-end distance of eukaryotic SR. **b-d**, Location of the FRET dyes are shown in the structures of RNC-bound SRP (**b**; PDB:3JAJ) and the RNC•SRP•SR complex in the distal conformation (**c, d**; PDB: 6FRK). The estimated distance between the dye pair is ~44 Å in (**b**), ~39 Å in (**c**), and ~36 Å in (**d**). **e-h**, smFRET histograms of signal sequence-fused SRP labeled with the Proximal (**e, f**) or Distal (**g, h**) probes without (**e, g**) and with (**f, h**) SR present. Asterisk indicates the fluorescently-labeled species. ‘pdf’, probability density function; E*, uncorrected FRET efficiency. ‘n’, the number of bursts used to construct each histogram, obtained from at least three independent measurements. The data were fit to the sum (solid line) of three Gaussian functions low, medium, and high FRET (dotted lines). The fractions of each population are indicated. **i-j**, smFRET histograms of SR labeled with Compaction probes in the absence (**i**) and presence (**j**) of signal sequence-fused SRP. These histograms were not fit, because the intermediate E* values arise from dynamic sampling of SR rather than discrete conformational states (Fig. 2n).

However, multiple challenges remain for initiation of protein translocation after assembly of the targeting complex. First, SRP and the Sec61p translocase share extensively overlapping binding sites at the ribosome exit site and on the nascent polypeptide^12,13^. Given this overlap, how RNCs are transferred from SRP to Sec61p is a long-standing puzzle. Second, eukaryotic SR is an α/β heterodimer anchored at the ER via association of the SRα X-domain with SRβ, an integral membrane protein^14,15^. A ~200 amino acid disordered linker separates the SR NG-domain from its membrane proximal X/β domains^16,17^, potentially posing another barrier for cargo loading onto the membrane-embedded Sec61p. Finally, activated GTP hydrolysis in the SRP•SR complex acts as a double-edged sword: while GTP hydrolysis drives disassembly of SRP from SR for their turnover^18,19^ and is not required for protein translocation *per se*^18,20^, premature GTP hydrolysis could abort targeting before the RNC engages Sec61p and/or other translocases^21,22^. Whether and how the timing of GTP hydrolysis is regulated in the mammalian SRP pathway is unclear.

Early cryo-electron microscopy (EM) analyses observed a loss of density for SRP54-NG at the ribosome exit site upon SR addition^11,23^. A recent cryo-EM structure of the RNC•SRP•SR complex showed that SRP can adopt a distinct conformation in which its NG-domain, bound to SR, docks at a ‘Distal’ site at the opposite end of 7SL RNA where SRP68/72 is located (Fig. 1a, middle and Fig. 1c)^23,24^. These observations suggest that SRP is dynamic and undergoes largescale conformational rearrangements after assembly with SR. However, these rearrangements have not been directly observed, and no information is available about their mechanism, regulation, and function.

To address these questions, we studied global conformational changes in human SRP and SR using Förster Resonance Energy Transfer (FRET). We developed three FRET pairs that monitor distinct molecular movements in the targeting complex. Detachment of SRP54-NG from the ribosome exit was detected using a donor dye (Atto550) labeled at SRP19(C64) and an acceptor dye (Atto647N) labeled at SRP54(C12) (Fig. 1a and b, Proximal Probes or SRP_Prox_)^5,9^. Docking of SRP54-NG at the distal site was detected using a donor dye (Cy3B) labeled at SRP54(C47) and Atto647N labeled near SRP68(P149) (Fig. 1a and c, Distal Probes or SRP_Dist_). Finally, we monitored the end-to-end distance of SR using Atto550 labeled at the C-terminus of SRαNG and Atto647N labeled at the N-terminus of SRβ (Fig. 1a and d, Compaction Probes or SR_Comp_). We detected FRET between all three pairs of probes using a diffusion-based single molecule technique with microsecond timescale Alternating Laser Excitation (μs-ALEX)^25–27^ (Extended Data Fig. 1). Unless otherwise specified, all measurements were made in the complete targeting complex in which SRP•SR is also bound to the ribosome and signal sequence and used a soluble SR complex in which the SRβ TMD, dispensable for SRP binding and cotranslational protein translocation, is removed^28^. Fluorescently labeled SRP and SR retain the ability to target preproteins to the ER (Extended Data Fig. 3 and Lee JH et al.^5^). We confirmed that the tested reaction conditions did not alter photo-physical properties of fluorophores, so that the observed FRET changes can be ascribed solely to conformational changes in SRP and SR (Extended Data Fig. 2).

Signal sequence- and ribosome-bound SRP_Prox_ mainly displayed a high FRET population (Fig. 1e), indicating that SRP54-NG initially docks near the ribosome exit site as previously reported^5^. In contrast, ~70% of SRP_Prox_ displayed low FRET upon SR addition (Fig. 1f), indicating that interaction with SR induces SRP54-NG to move away from the ribosome exit. The opposite was observed with the distal probes: the FRET histogram of SRP_Dist_ was dominated by low FRET populations (Fig. 1g), whereas approximately 34% of the complex acquired high FRET upon addition of SR (Fig. 1h), indicating acquirement of the distal state in this population. These results provide direct evidence that the SRP-SR NG-complex detaches from the ribosome exit site, where it initially assembles^5^, and docks at the distal site where SRP68/72 is located. Intriguingly, the population of the targeting complex that exhibited low FRET detected by the proximal probes far exceeded the high FRET population detected by the distal probes, suggesting that the targeting complex samples additional conformations in which the NG-complex is not stably docked at either the ribosome exit or the distal site.

Finally, we monitored the global conformational changes of SR using the compaction probes that report on the proximity of its folded NG- and X/β-domains (Fig. 1a and d). As SR was implicated in ribosome binding^16,17^, we first measured the conformation of free SR with the 80S ribosome present. The smFRET histogram of SR_Comp_ exhibited a main peak at FRET ~ 0.15 (Fig. 1i), indicating that the NG- and X/b domains are separated by ≥90 Å in free SR. When signal sequence- and ribosome-bound SRP were present, however, the FRET distribution of SR_Comp_ became broader and shifted to higher FRET with a major peak at FRET ~ 0.7 (Fig. 1j). These results show that SR undergoes a global compaction upon binding with cargo-loaded SRP, bringing its NG-domain much closer to the membrane-proximal X/β-domain.

To understand the molecular mechanisms that drive these conformational changes, we introduced mutations that disrupt the interaction surfaces of SRαNG with SRβ, SRX, or SRP68/72 (Fig. 2a, Extended Data Fig. 4a and 4b). Additionally, we characterized one of the SRP54 mutations (G226E) that cause congenital neutropenia with Shwachman-Diamond-like features^7,8^ (Fig. 2a and Extended Data Fig. 4a). None of the mutations impaired SRP-SR complex assembly or their reciprocal GTPase activation (Extended Data Fig. 4c-e). As efficient SRP-SR interaction requires both the ribosome and signal sequence^5^, these results also ruled out defects of these mutants in ribosome binding or signal sequence interaction. Thus, all of the defects observed in the following analyses are caused by conformational defects that occur after SRP-SR assembly.

**Figure 2.**
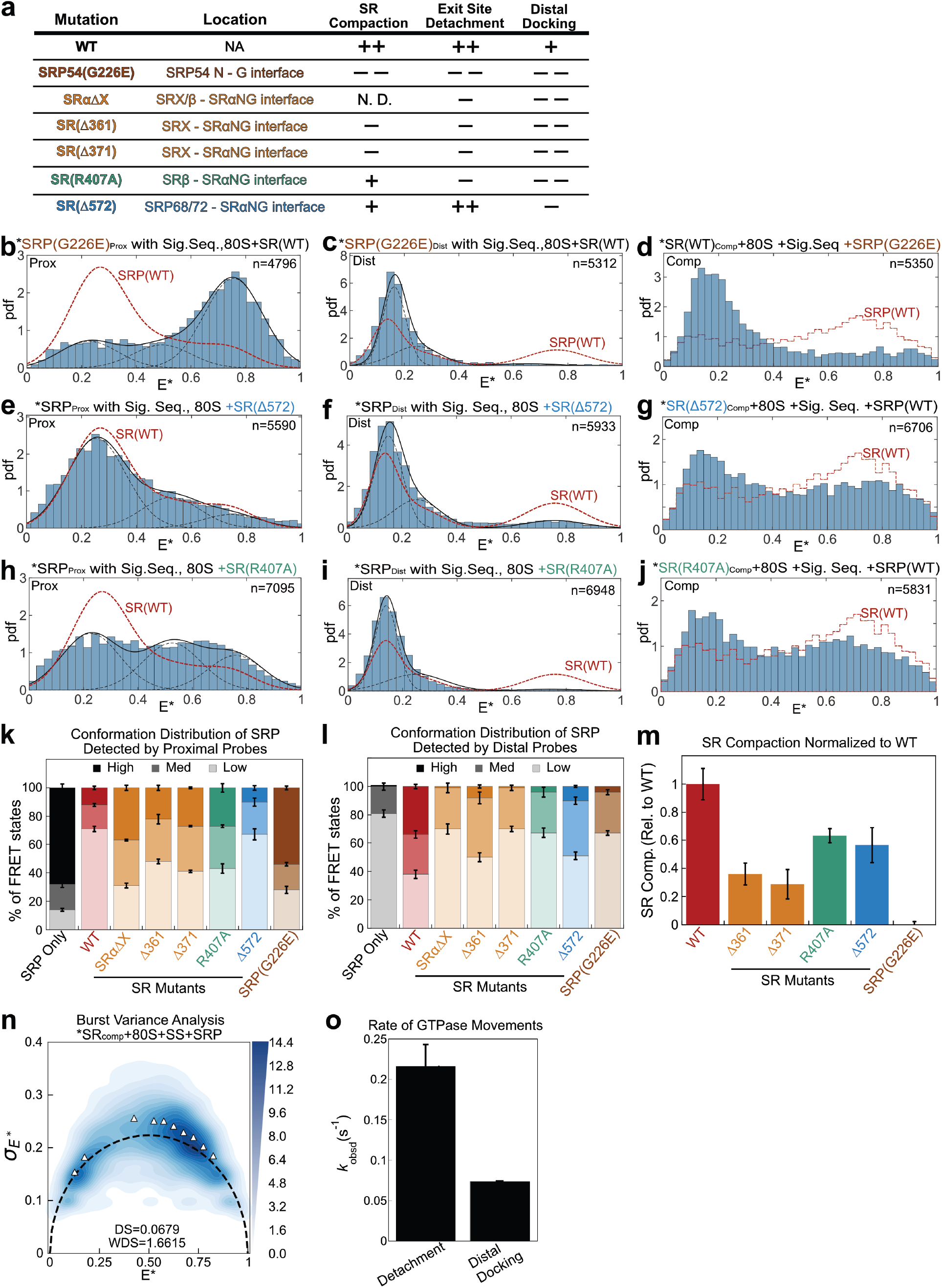
SR compaction drives GTPase movements in the targeting complex. **a**, Summary of the mutations characterized in this study and their phenotypes derived from the data in **b-m**. Mutations are colored based on the interactions disrupted. Details of each mutation are shown in Extended Data Fig. 4a-b. **b-j**, smFRET histograms of targeting complex containing mutants SRP(G226E) (**b-d**), SR(D572) (**e-g**), or SR(R407A) (**h-j**) detected by the Proximal (**b, e, h**), Distal (**c, f, i**), and Compaction (**e, g, j**) probes. The data were analyzed as in Figure 1. The red dotted lines outline the corresponding histograms of the wild-type targeting complex. **k-l**, Quantification of the populations in low 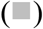, median 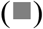, and high 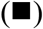 FRET states detected by the Proximal (**k**) or Distal (**l**) probes. **m**, Quantification of SR compaction, calculated from the fraction of targeting complex displaying high FRET (E* = 0.6 – 0.8) and subtracting the corresponding value in the histogram of ribosome-bound SR. All values are normalized to that of the wildtype targeting complex. Error bars in **k-m** denote SD from at least three independent experiments. **n**, BVA plot of SR compaction in the targeting complex. The black dashed curve depicts the static limit. Triangles denote the observed standard deviation for individual E* bins (*SDE\**) and were used to calculate the dynamic score (DS) and weighted dynamic score (WDS) (eqs 9-11 in Supplementary Methods). o, Rate constants for detachment of the NG-complex from the ribosome exit (0.21 ± 0.027 s^-1^) and for distal site docking (0.07 ± 0.0007 s^-1^) measured using the Proximal and Distal probes, respectively. Rate constants are from the data in Extended Data Fig. 6d and are shown as mean ± SD, with n = 3-5.

ALEX measurements suggest that these mutants block the conformational rearrangements in the targeting complex at distinct steps. SRP54(G226E), which causes congenital neutropenia^7,8^, severely impaired all three conformational rearrangements in the targeting complex (Figs. 2b-d; summarized in Fig. 2k-m, *brown*). Similar albeit less pronounced defects were observed with mutants SR(D361) and SR(Δ371) that disrupt the intramolecular interactions between SRX and SRαNG: they not only compromised SR compaction, as expected (Extended Data Fig. 5c and 5f; summarized in Fig. 2m, *orange*), but also impaired the detachment of the NG-domain complex from the ribosome exit site and its docking at the distal site (Extended Data Fig. 5a, b, d and e; summarized in Figs. 2k and 2l, *orange*). These results suggest that the intramolecular interactions within SR are crucial for the movement of the NG-complex from the proximal to the distal site. Reciprocally, all the mutations that disrupted distal site docking also reduced SR compaction to varying degrees (Fig. 2m), suggesting that the distal state stabilizes a highly compact SR. Nevertheless, several mutants showed more specific defects. SR(Δ572), which disrupts the contact of SRαNG with SRP68/72 (Figs. 2a, Extended Data Fig. 4a and b, *blue*), specifically destabilized distal site docking but did not affect the removal of the NG-domain complex from the ribosome exit and modestly reduced SR compaction (Fig. 2e-g; summarized in Fig. 2k-m, *blue*). This shows that distal docking is not required for, and probably occurs after, the other rearrangements. Finally, SR(R407A) disrupted the interaction of SRαNG with SRβ (Figs. 2a and Extended Data Fig. 4a, 4b, 6e, *green*). This mutant was impaired in both of the lateral movements of the NG-domain complex as strongly as SR(D361) and SR(D371) (Fig. 2h and 2i; summarized in Fig. 2k and 2l, *green*) but was able to undergo significant compaction similarly to SR((D572) (Fig. 2j and 2m, *green*), suggesting that SR can sample the compact conformation before the other rearrangements. The distinct mutational phenotypes (qualitatively summarized in Fig. 2a) suggest a sequential model in which SR compaction precedes and potentially drives the movement of the NG-complex from the ribosome exit to the distal site of SRP.

Analysis of the dynamics of the rearrangements supported this sequential model. We first performed burst variance analysis (BVA), which detects dynamics by comparing the standard deviation of E* (σ_E*_) for individual molecules to the static limit, defined by photon statistics (Supplementary Methods)^27,29–31^. If multiple conformations interconvert on the sub-millisecond timescale, the observed sE* would be higher than the static limit, whereas σ_E*_ would lie on the static limit curve if conformational interconversions are slower compared to molecular diffusion (1-5 milliseconds)^27,29–31^. SR_Comp_ displayed σ_E*_ values significantly higher than the static limit (Fig. 2n, triangles versus dashed curve). This indicates that SR samples the compact state on the submillisecond timescale and is consistent with the disordered nature of the SR linker^16,17^. In contrast, σ_E*_ for SRP_Prox_ and SRP_Dist_ are much closer to the static limit (Extended Data Fig. 6a-c). Kinetic measurements using the SRP_Prox_ and SRP_Dist_ probes further showed that exit site detachment and distal docking occur with rate constants of 0.21 s^-1^ and 0.07 s^-1^, respectively (Fig. 2o and Extended Data Fig. 6d). Thus, SR can rapidly undergo compaction upon binding of cargo-loaded SRP, followed by detachment of the NG-complex from the ribosome exit and subsequent docking at the distal site.

Structural modeling supported SR compaction as a driver for the GTPase movements. When the compacted SRP54-NG•SR structure was superimposed on SRP54-NG bound at the ribosome exit site^9,24^, we found that SR compaction brings SRX close to the ribosome, potentially generating a steric clash that would destabilize the proximal state and drive detachment of the NG-complex from the ribosome exit (Extended Data Fig. 6e). This is consistent with smFRET data showing that all the SR mutations that impair this detachment disrupt interactions within the SR complex (D361, D371, and R407A). An alternative model consistent with most of the data could involve initial formation of bidentate interactions of SRP with an extended SR via both the NG-domain contacts at the ribosome exit and SRX/b docking at the distal site, followed by SR compaction that moves the NG-complex to the distal site. Intriguingly, SRP54(G226E), which caused the most severe defects in all three rearrangements, is located at the interface between the N- and G-domains of SRP54 (Extended Data Fig. 6, *brown*). In the extensively-studied bacterial SRP pathway, this interface acts as a fulcrum that undergoes cooperative adjustments in both the SRP54 and SR NG-domains upon their GTP-dependent assembly^32–34^. These observations suggest that the cooperative rearrangements within the SRP•SR NG-domain complex during its assembly are coupled to restructuring of the SR linker and thus amplified into the largescale molecular motions observed here.

We next asked how the conformational changes in SRP and SR are regulated by their biological cues including the ribosome and signal sequence. The smFRET histogram of apo-SR_Comp_ had a broad distribution without a major peak (Fig. 3a), indicating that free SR samples multiple conformations. With either the ribosome (Fig. 1i and 3a-d, dashed red line) or SRP (Fig. 3b) present, the histogram was dominated by a peak at FRET < 0.2, indicating that binding of the ribosome or SRP induced SR into a highly extended conformation. However, when both the ribosome and SRP are present, the histogram for SR_Comp_ peaked at FRET ~ 0.7 (Fig. 3d) and was similar to that in the presence of ribosome- and signal sequence-bound SRP (Fig. 1j). Signal sequence-bound SRP also shifted the histogram of SR_Comp_ to higher FRET, but less effectively than ribosome-bound SRP (Fig. 3c). Thus, the ribosome and SRP cooperatively drive the compaction of SR.

**Figure 3.**
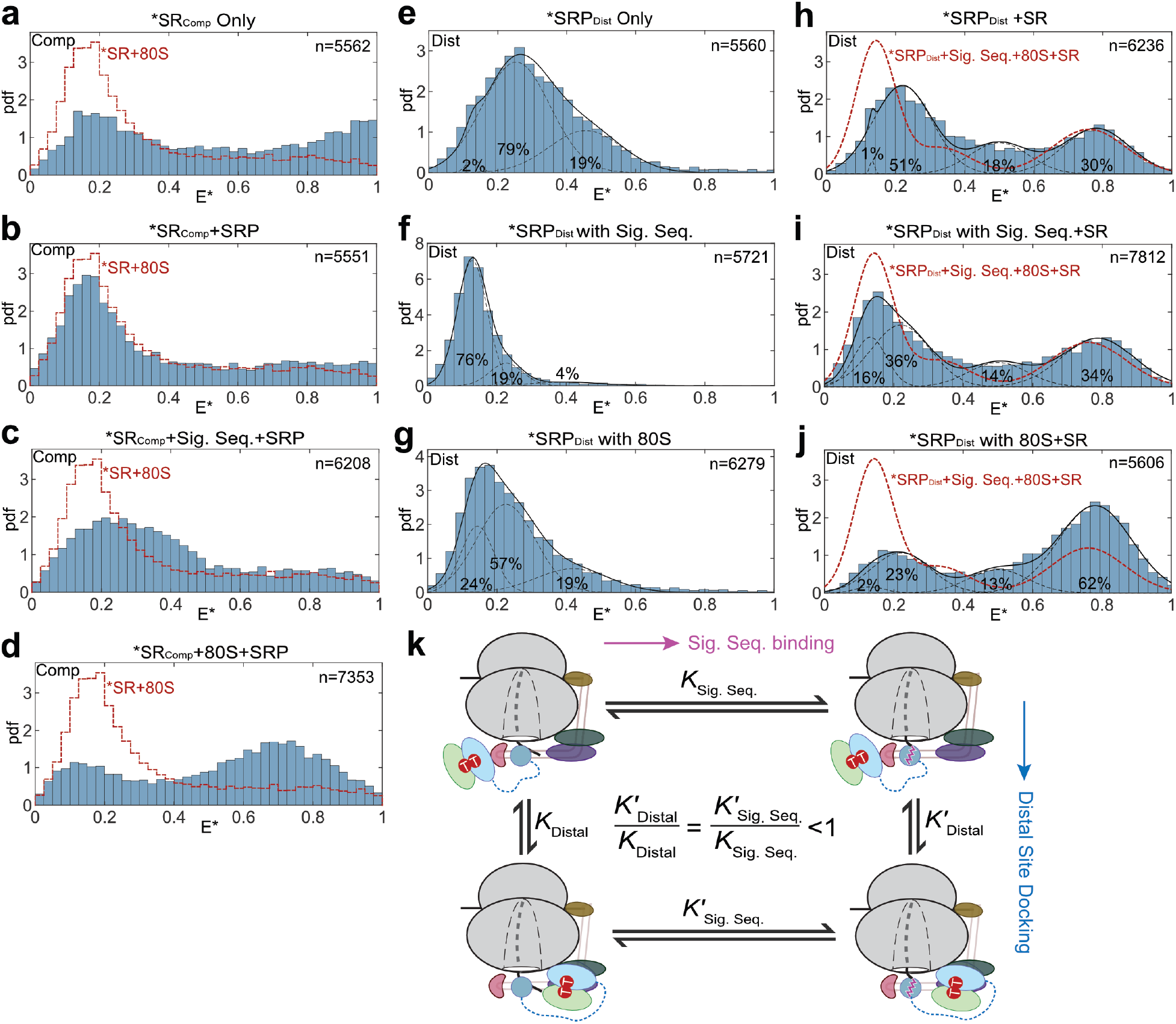
The ribosome and signal sequence regulate conformational changes in the targeting complex. **a-d**, smFRET histograms of SR_Comp_ in the absence (**a**) or presence of SRP (**b**), signal sequence bound SRP (**c**), or ribosome-bound SRP (**d**). The red dotted lines outline the data for ribosome-bound SR_Comp_. **e-j**, smFRET histograms of SRP_Dist_ (**e-g**) or the SRP_Dist_oSR complex (**h-j**) in the absence (**e, h**) or presence of signal sequence (**f, i**) or ribosome (**g, j**). The red dotted lines in **h-j** outline the data with the complete targeting complex. **k**, Thermodynamic cycle analysis of the coupled equilibria of signal sequence binding and distal site docking of the NG-complex. The less favorable distal site docking in the presence of signal sequence (*K*’_Dist_ < *K*_Dist_) implies weaker signal sequence binding in the distal state (*K*’_Sig. Seq_. < *K*_Sig. Seq_).

We further tested how the lateral movements of the GTPase complex are regulated. We previously showed that the histogram of free SRP_Prox_ is dominated by a medium FRET population; signal sequence drives SRP to the proximal state and thus shifts the histogram toward higher FRET, whereas the ribosome induces SRP to sample at least three conformations (Lee JH *et al.*^5^ and Extended Data Fig. 7a-c). The smFRET histograms of SRP_Dist_ largely mirrored these ribosome- and signal sequence-induced changes in SRP and indicated the absence of the distal state prior to SR binding (Fig. 3e-g). Addition of SR induced the distal state at FRET ~ 0.8 under all tested conditions (Fig. 3h-j). Surprisingly, in the presence of ribosome, the distal state was nearly two-fold more enriched (62%) than in the complete targeting complex with signal sequence present (Fig. 3j vs 3i). These results show that distal docking of the GTPase complex is driven by SR and favored by the ribosome, but is destabilized by the signal sequence.

Changes in the equilibrium for distal site docking with and without signal sequence is thermodynamically coupled to the changes in signal sequence interactions before and after distal docking (Fig. 3k). Therefore, the less favorable distal docking in the presence of signal sequence (*K*’_Distal_ <*K*_Distal_) implies that the interaction of SRP with signal sequence is destabilized upon rearrangement to the distal state (*K*’_Sig. Seq_. < *K*_Sig. Seq_; Fig. 3k). This is consistent with the weaker EM density of the signal sequence in the distal state structure compared to the RNC•SRP complex^9,24^, and suggests that distal site docking provides a mechanism to facilitate the handover of the nascent polypeptide from SRP to the translocation machinery.

GTP hydrolysis in the SRP•SR complex drives their irreversible disassembly and is an important regulatory point in the bacterial SRP pathway^35,36^. To test how the conformational rearrangements in the SRP•SR complex regulate GTP hydrolysis, we measured the stimulated GTPase activity of the targeting complex (*k*_cat_) using mutants and conditions that bias the conformational equilibria (Extended Data Fig. 4c and 4d)^5^. The complete targeting complexes assembled with all the SR conformational mutants displayed higher GTPase rates (*k*_cat_) than the wildtype complex (Fig. 4a). The value of *k*_cat_ negatively correlated with the fraction of SRP•SR in the distal state (Fig. 4b). The lowest GTPase rate was observed with ribosome-bound SRP•SR^5^, which strongly favors the distal state (Figs. 4b and 3j). This strongly suggests that docking at the distal site inhibits GTP hydrolysis and thus increases the lifetime of the targeting complex at the ER membrane. This is consistent with our previous observation that mutations in the SRP72 C-terminus, which is positioned near the GTPase active site in the distal state structure, hyper-activated the GTPase reaction^24^. On the other hand, the targeting complex bearing mutant SRP(G226E), in which the NG-domain complex is trapped at the ribosome exit site, hydrolyzed GTP at a rate constant of ~5 min^-1^, providing an estimate for the GTPase rate when the targeting complex is in the Proximal state. These observations further suggest that hyperactive GTP hydrolysis in the targeting complex primarily occurs when the NG-complex is not docked at either the ribosome exit or the distal site. Thus, conformational rearrangements tune the timing of GTP hydrolysis, possibly providing a balance between efficient SRP turnover and substrate transfer.

**Figure 4.**
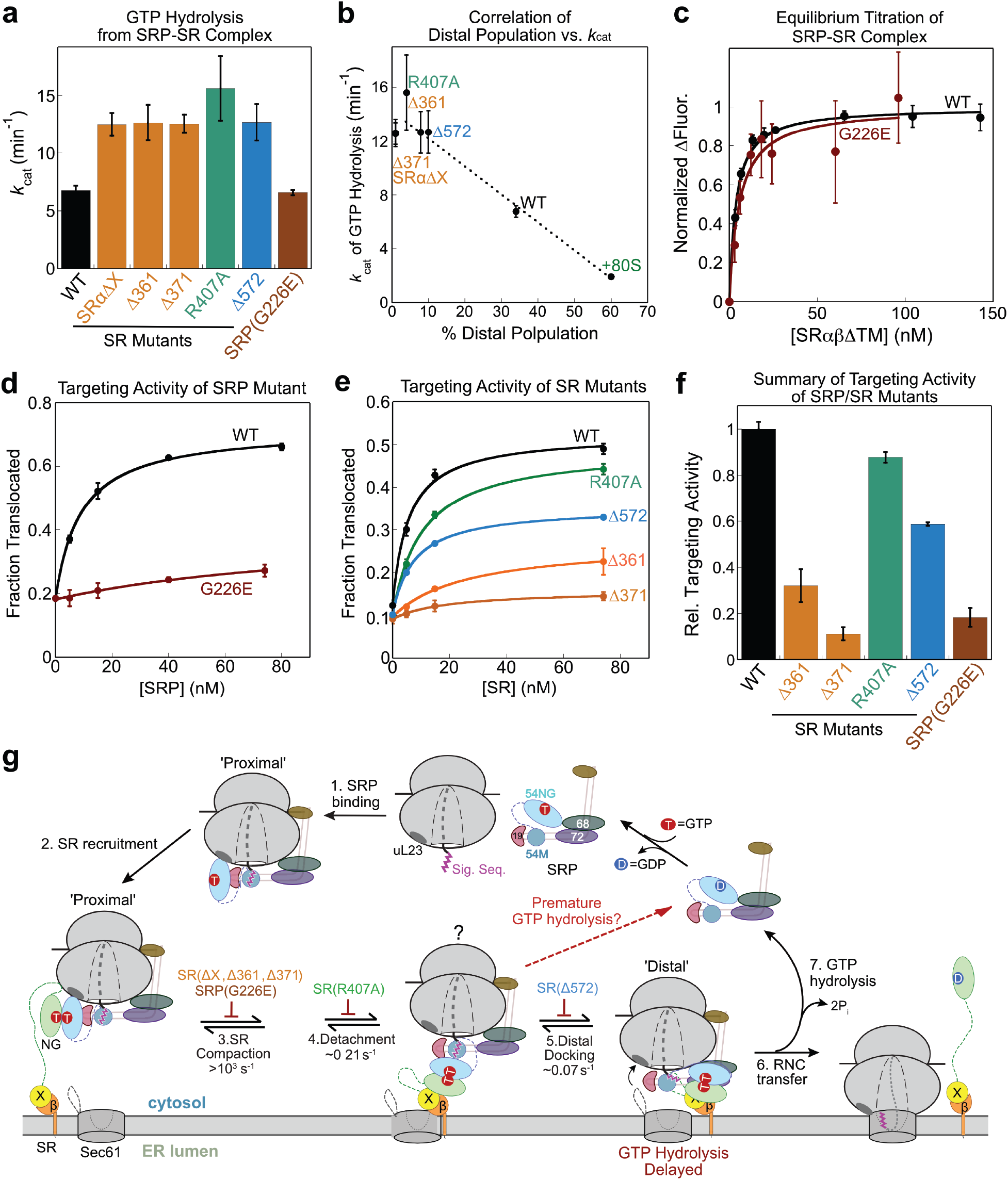
Post-targeting conformational changes in SRP and SR are essential for protein translocation. **a**, Summary of the rate constants of stimulated GTP hydrolysis (*k*_cat_) in targeting complexes assembled with wildtype protein or the indicated SRP and SR mutants. Values are reported as mean ± SD, with n = 3. **b**, *k*_cat_ negatively correlates with the fraction of SRP•SR complex in the distal conformation. The *k*_cat_ for the reaction with ribosome-bound SRP•SR complex (‘+80S’) was measured previously^5^. **c**, Equilibrium titrations to measure the binding affinity between SRP(G226E) and SR. The data were fit to Eq 4 and gave equilibrium dissociation constants (*K*_d_) of 3.7 ± 0.35 and 5.6 ± 1.7 nM for wildtype SRP and mutant SRP(G226E). Error bars represent SD, with n = 3. **d-e**, Co-translational translocation of pPL mediated by wildtype or the indicated SRP (**d**) and SR (**e**) mutants. Translocation efficiencies were quantified from the data in Extended Data Fig. 8a, b and are reported as mean ± SD, with n = 2-3. **f**, Summary of the translocation efficiency of each mutant relative to the wildtype targeting complex at saturating protein concentrations. **g**, Model for co-translational protein targeting and translocation. SRP is pre-organized into the Proximal conformation upon binding to signal sequence bearing ribosomes (step 1) and recruits SR via interaction between their NG-domains (step 2). This induces SR compaction (step 3) to drive the detachment of the NG-complex from the ribosome exit (step 4). The question mark on the targeting complex after step 4 denotes that the precise structure of this intermediate is unknown. Docking of the GTPase complex at the distal site (step 5) further primes the RNC for unloading to Sec61p (step 6). GTP hydrolysis, which dissociates SRP from SR (step 7), is delayed in the Distal state, and failure to reach the distal conformation could cause premature GTP hydrolysis that aborts targeting (dashed red arrows).

Finally, we tested the role of these conformational rearrangements in SRP function using a reconstituted assay that measures co-translational translocation of a model substrate, preprolactin (pPL), to ER microsomes^5,37^ mediated by the SRP and SR variants (Extended Data Fig. 3a). Most of the SR conformational mutants are defective in pPL translocation (Fig. 4e, 4f, and Extended Data Fig. 8a). The largest defects were observed with SR(D361) and SR(D371), which block the rearrangements at the earliest stage (Fig. 2, Fig. 4e, and 4f, *orange*). SR(D572), which specifically blocks distal site docking, also substantially reduced translocation efficiency (Fig. 2, Fig. 4e and 4f, *blue*), supporting a role of the distal state in ensuring efficient protein translocation. The only exception was SR(R407A), which did not substantially affect pPL translocation despite impairments in the lateral movements of the NG-complex; this might reflect contributions from additional factors in the cell lysate and ER microsomes during translocation measurements that were not present in smFRET measurements of the purified targeting complex. Notably, mutant SRP(G226E), which causes severe congenital neutropenia^7,8^, strongly impaired pPL translocation (Fig. 4d and 4f, *brown*). Contrary to a previous report^7^, we found that SRP(G226E) displays basal GTPase activity and stimulated GTPase reactions with SR as efficiently as wildtype SRP (Extended Data Fig. 4d and 4f). Using a pair of FRET probes incorporated at SRP54(C47) and the SRα C-terminus^5^, equilibrium titrations further corroborated that SRP(G226E) assembles a stable complex with SR (Fig. 4c), showing that this mutant was specifically blocked in conformational rearrangements after SRP-SR assembly (Fig. 2b-d). These results demonstrate that the post-targeting conformational rearrangements in SRP and SR play essential roles in co-translational protein translocation and can be the point of failure in devastating pathology.

The results here demonstrate that, after an RNC•SRP•SR complex assembles at the ER, multiple largescale conformational rearrangements in the targeting complex are required to initiate protein translocation (Fig. 4g). During cargo recognition, signal sequence bearing ribosomes induce SRP into a ‘Proximal’ conformation in which the SRP54-NG domain docks at uL23 near the ribosome exit (step 1)^5^. In this conformation, SRP rapidly recruits SR via interaction between their NG-domains (step 2). Cooperative rearrangements in the NG-complex upon its assembly, especially those at the N-G domain interface, are sensed by the SR linker and amplified into a global compaction of SR, likely bringing the translating ribosome near the ER membrane (step 3). This compaction also drives the detachment of the GTPase complex from the ribosome exit (step 4), exposing universal docking sites at the ribosome exit site for subsequent interaction with the Sec61p translocase. A fraction of the GTPase complex docks at the distal site (step 5). In this state, the interaction of SRP with the signal sequence is destabilized to further prime the RNC for subsequent unloading, and delayed GTP hydrolysis could generate an extended time window during which the targeting complex searches for and allows the translating ribosome to engage the appropriate translocase (step 6). These post-targeting molecular movements resolve multiple mechanistic challenges during initiation of protein translocation and are demonstrable points of failure in diseases such as congenital neutropenia, and could serve as important points for biological regulation as well as disease intervention.

## Methods

### Vectors

The vectors for expression of SRP and SR subunits and for fluorescence labeling of SRP19, SRP54 and SRα C-terminus have been described^5^. To fluorescently label SRP68, an Sfp recognition motif (ybbr6, DSLEFI)^38^ was inserted after Pro149 using Fastcloning. To fluorescently label SRβΔTM, a longer Sfp recognition motif (ybbr11, DSLEFIASKLA)^38^ was inserted at the SRβ N-terminus using Fastcloning. Expression vectors for mutant SRP and SRs were generated using QuickChange mutagenesis protocol (Stratagene).

### Biochemical Preparations

Wildtype and mutant SRP and SR proteins were expressed and purified as described^5^. Mammalian SRP was prepared as described^5^. Briefly, SRP protein subunits were expressed and purified in bacteria or yeast. A circularly permutated 7SL RNA variant was *in vitro* transcribed and purified on a denaturing polyacrylamide gel. SRP was assembled by first refolding 7SL RNA and sequentially adding SRP19, SRP68/72, SRP9/14, and SRP54. Holo-SRP was purified using a DEAE ion-exchange column. Unless otherwise specified, the C-terminus of human SRP54 was fused to the 4A10L signal sequence (hSRP54-4A10L) to generate signal sequence-bound SRP, as described^5^. Ribosome from rabbit reticulocyte lysate (RRL) was purified by ultracentrifugation through a sucrose gradient, as described^5^. The use of ribosome and signal sequence fusion to SRP54 reproduced the effects of signal sequence-bearing RNCs on the conformation and activity of SRP^5^.

### Fluorescence Labeling

SRP54(C12), SRP54(C47), and SRP19(C64) were labeled with Atto550, Atto647N, or Cy3B using maleimide chemistry as described^5^.

SRP68/72 was labeled via Sfp-mediated conjugation of CoA-Atto647N or CoA-Cy3B at Ser2 in ybbr6 tagged SRP68 following the procedure described in ^38^. The labeling reaction contained 0.4 molar ratio of protein to Sfp enzyme and a 3-fold excess of CoA-Atto647N (or CoA-Cy3B), and was carried out for 20 minutes at room temperature in Sfp-labeling Buffer (50 mM KHEPES (7.5), 10 mM MgCl_2_, 150 mM NaCl, and 20% glycerol). Labeling efficiency was close to 100%. Labeled SRP68/72 was immediately used for SRP assembly.

SRαβΔTM was doubly labeled via sortase-mediated ligation at the SRα C-terminus and Sfp-mediated conjugation of CoA-dye at the N-terminal ybbr11 tag on SRβ. The labeling reaction contained 0.4 molar ratio of Sfp to protein and a 2-fold excess of CoA-Atto647N (or CoA-Atto550), and was carried out for 30 minutes at room temperature in Sfp-labeling Buffer. A 4-fold molar excess of sortase,10-fold excess of GGGC-Atto550 (or GGGC-Atto647N), and 0.1 volume of 10X Sortase Buffer (500mM Tris-HCl 7.5, 1.5M NaCl, and 100 mM CaCl2) was then added, and the labeling reaction was carried out for an additional 3 hours at room temperature. Labeled SRP68/72 was purified using Ni-Sepharose resin. Labeling efficiency was close to 100% for the Sfp reaction and ~60–70% for the sortase reaction.

### Biochemical Assays

All proteins except for SRP were ultracentrifuged at 4 °C, 100,000 rpm in a TLA100 rotor for 30 – 60 minutes to remove aggregates before all assays. GTPase reactions were performed in SRP Assay Buffer (50 mM KHEPES (pH 7.5), 150 mM KOAc, 5 mM Mg(OAc)_2_, 10% glycerol, 2 mM DTT, and 0.04% Nikkol) at 25 °C and were followed and analyzed as described^5,39^. Details for the determination of the GTPase rate constants are described in Supplementary Methods. Co-translational targeting and translocation of pPL into salt-washed and trypsin digested rough ER microsomes (TKRM) were performed and analyzed as described in Supplementary Methods. Steady-state fluorescence measurements were carried out on a Fluorolog 3-22 spectrofluorometer (Jobin Yvon) at 25 °C in SRP Assay Buffer. Acquisition and analyses of fluorescence data are described in Supplementary Methods.

### smFRET

Measurements were performed as described^5,25,26^. Labeled SRP was diluted to 100–200 pM in SRP Assay Buffer containing 200 μM non-hydrolysable GTP (GppNHp), 150 nM 80S, and 1.5 μM SRαβΔTM where indicated. To measure the conformation of SR, doubly labeled SRαβΔTM was diluted to 100-200 pM in SRP Assay Buffer containing 200 μM GppNHp, 400 nM 80S, and 400 nM SRP or SRP-4A10L where indicated. Data were collected over 30-60 min using an ALEX-FAMS setup with two single-photon Avalanche photodiodes (Perkin Elmer) and 532 nm (CNI laser) and 638 nm (Opto Engine LLC) continuous wave lasers operating at 150 μW and 70 μW, respectively. Analysis of μs-ALEX data are described in Supplementary Methods.

## ACKNOWLEDGMENT

We thank S. Chandrasekar and H. Hsieh for sharing reagents, A. Jomaa and N. Ban for helpful discussions, and members of the Shan lab for comments on the manuscript. This work was supported by National Institutes of Health grant GM078024 and the Gordon and Betty Moore Foundation through grant GBMF2939 to S.-o. Shan, and by National Institutes of Health grant GM130942 and Dean Willard Chair funds to S.W.

## AUTHOR CONTRIBUTIONS

J.L., Y.H., and S.S. designed research; J.L., Y.H., R.Q, and X.S. performed biochemical experiments and analyzed data; J.L., R.Q., and S.C. performed μs-ALEX experiments and analyzed data; S.W. provided guidance for μs-ALEX analysis; J.L. and S.S. wrote the manuscript with input from S.C. and S.W.

