## Extended Data Figures for "Receptor compaction and GTPase movements drive cotranslational protein translocation"

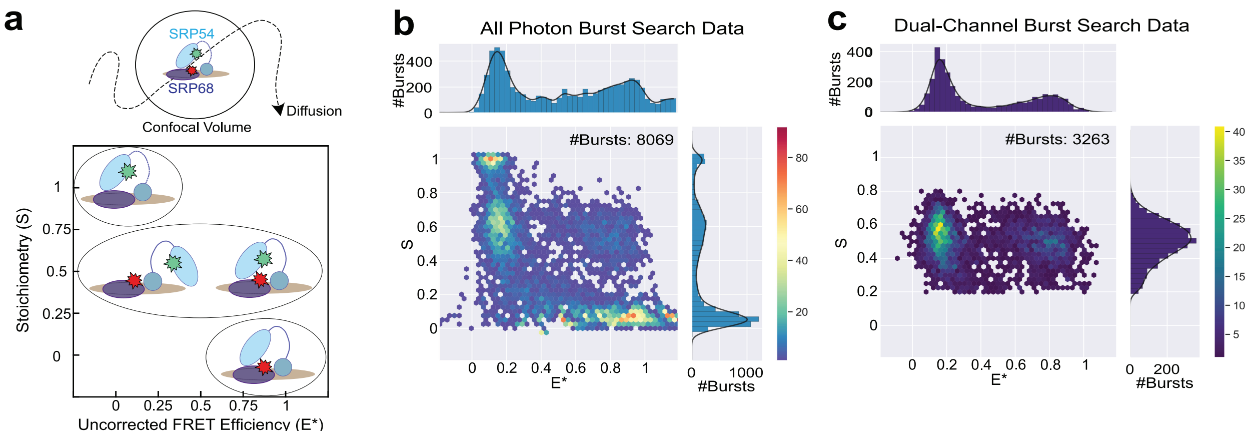


**Extended Data Figure 1.** Diffusion-based smFRET experiments. **a,** Schematic of the fluorescence-aided molecular sorting measurements using microsecond timescale Alternating Laser Excitation Spectroscopy (μs-ALEX). Fluorescently labeled SRPs diffusing through a femtoliter-scale confocal volume are alternately excited with donor (green) and acceptor (red) excitation lasers. The fluorescence from both donor and acceptor dyes are measured to calculate donor-acceptor stoichiometry (S) and uncorrected FRET efficiency (E*) to generate 2-D E*-S plots. This allows optical purification of the doubly labeled particles that contain both donor and acceptor dyes (S ~ 0.5), and 1D-projection of the doubly labeled population onto the E* axis generates the FRET histogram[1–3]. **b-c,** 2-D E*-S plots of $SRP_{Dist}$ bound with signal sequence, ribosome, and SR generated by all photon burst search **(b)** and dual-channel burst search **(c).** The dual-channel burst search filters out singly labeled species such as donor only (S~1) and acceptor only (S~0).

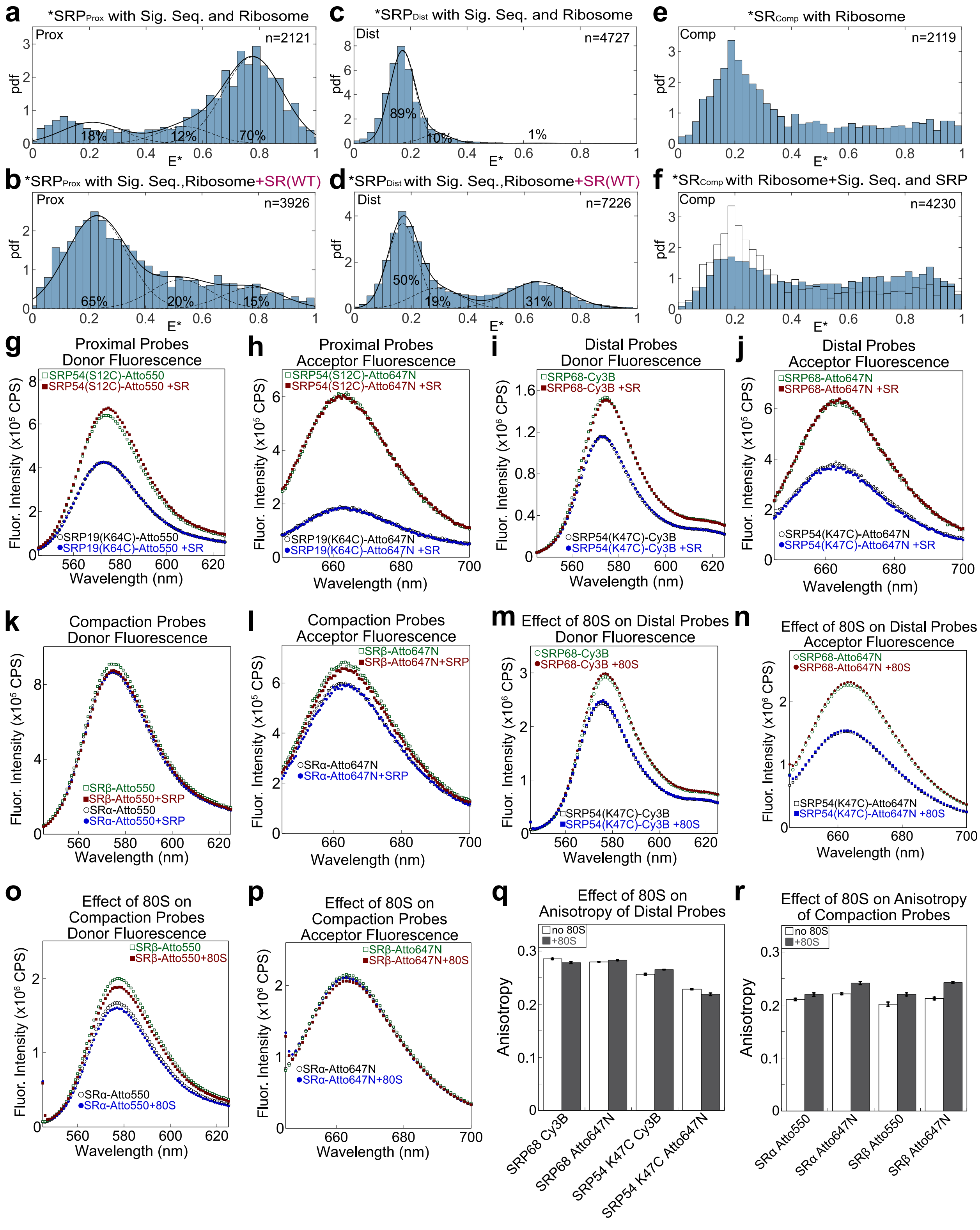


**Extended Data Figure 2.** The change in FRET efficiency is due only to the conformational change of the SRP. **a-d,** FRET histograms of $SRP_{Prox}$ (**a, b**) and $SRP_{Dist}$ (**c, d**) in the absence (**a, c**) or presence (**b, d**) of SR, with the position of the donor and accepter dyes switched compared to those in Figure 1. **e-f,** smFRET histograms of $SR_{Comp}$ bound to the ribosome with (**f**) and without (**e**) signal sequence-bound SRP present. The donor and acceptor dye positions were swapped compared to that in Figure 1. **g-l** Addition of SR or SRP did not affect the fluorescence intensity of the donor or acceptor dyes in $SRP_{Prox}$, $SRP_{Dist,}$ and $SR_{Comp}$. Steady-state fluorescence emission spectra are shown for SRP or SR singly labeled with the donor or acceptor dye at the indicated positions. Every labeling position was tested with both donor and acceptor dyes to exclude dye/position specific effects. The reactions contained the same concentrations of components as in the smFRET experiments, except that the labeled complexes were present at 15 nM. **m-p,** Addition of 80S did not affect the fluorescence intensity of the donor or acceptor dyes in $SRP_{Dist}$ and $SR_{Comp}$. Steady-state fluorescence emission spectra are shown for SRP or SR singly labeled with the donor or acceptor dyes at the indicated positions. The absence of an effect of 80S on the proximal probes have been reported previously[4]. **q-r,** Addition of 80S did not significantly affect the fluorescence anisotropy of the fluorescence dyes used for the Distal or Compaction probes. Fluorescence anisotropy was measured for the indicated fluorescence dyes labeled at the indicated positions in the absence and presence of 100 nM/400 nM 80S (for Distal/Compaction probes, respectively) to verify that the observed FRET changes are not due to dye orientational effects. Error bars represent SD, with n = 3.

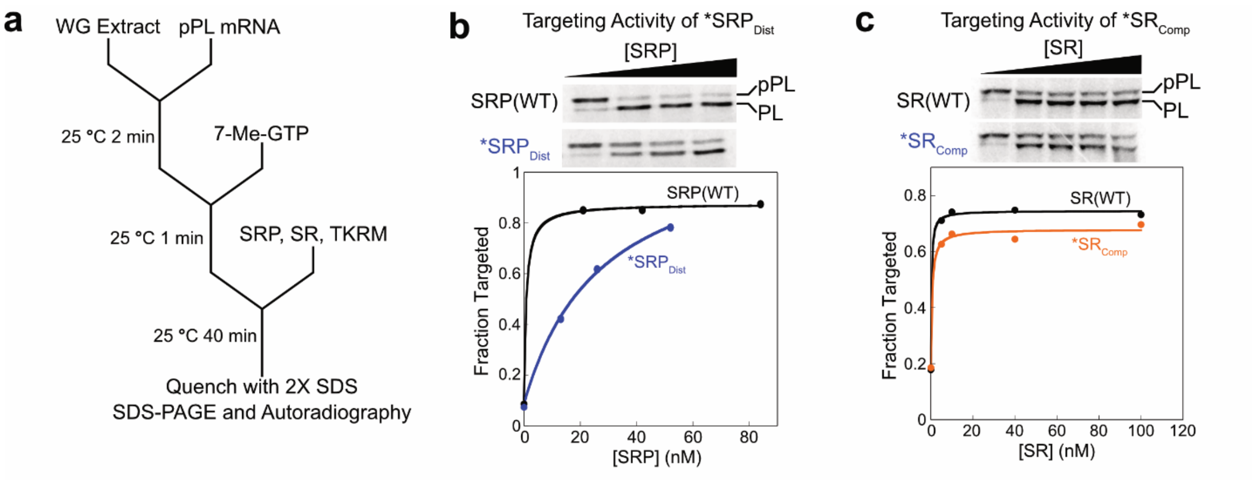


**Extended Data Figure 3**. Protein targeting and translocation activity of doubly-labeled SRP and SR. **a,** Schematic to depict the *in vitro* protein targeting assay. Purified mRNA of a model SRP substrate, preprolactin (pPL), was translated in wheat germ lysate containing $^{35}$S-methionine. 1.5 mM 7-methyl-GTP was added 2 minutes after initiation of translation to inhibit additional rounds of translation. SRP, SR and high-salt washed, trypsin-digested rough ER microsomes (TKRM) were added to initiate the targeting reaction. The reactions were incubated at 25 °C for 40 min and quenched in SDS buffer, and analyzed by SDS-PAGE followed by autoradiography[4]. **b,** Upper panel, representative SDS-PAGE autoradiography gels showing the translocation of pPL by wildtype SRP or $SRP_{Dist}$. Reactions contained 100 nM SR, indicated concentrations of SRP, and 0.5 eq of TKRM. Lower panel, quantification of pPL translocation efficiency from the data. **c,** Upper panel, representative SDS-PAGE autoradiography gels showing the translocation of pPL by wildtype SR or $SR_{Comp}$. The reactions contained 20 nM SRP and 0.5 eq of TKRM. Lower panel, quantification of pPL translocation efficiency from the data.

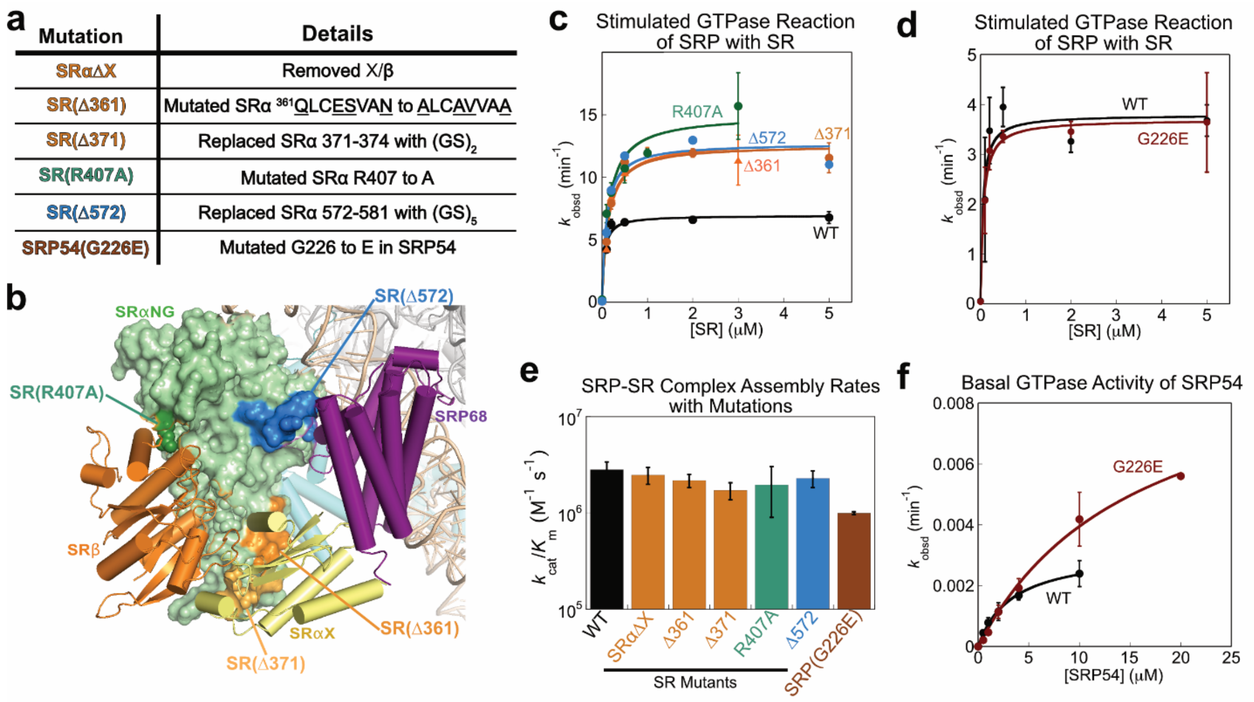


**Extended Data Figure 4**. Additional data for SRP and SR mutations that disrupt conformational changes in the targeting complex. **a,** Details for the mutations characterized in this study. **b,** The location of individual SR mutations are highlighted in the structure of the distal state complex (PDB: 6FRK)[5]. **c,** SR concentration dependences of the stimulated GTPase reaction between SRP and SR mutants. All reactions contained 0.2 μM SRP-4A10L,100 μM GTP, 0.25 μM 80S, and the indicated concentrations of wildtype or mutant SR. **d,** SR concentration dependences of the stimulated GTPase reaction between SRP(G226E) and SR. The reactions contained 0.2 μM SRP-4A10L or SRP(G226E)-4A10L, 500 μM GTP, and 0.25 μM 80S. The lines are fits of the data to Eq. 2 and the obtained rate constants ($k_{cat}/K_m$ and $k_{cat}$) are summarized in panel e and Fig. 4b, respectively. **e,** Summary of the values of $k_{cat}/K_m$ from parts (c) and (d). **f,** Basal GTPase reactions of wild-type SRP and SRP(G226E). Reactions were measured as described in Supplementary Methods. The lines are fits of the data to Eq. 1 and gave rate constants of $k_{cat,wt}$ = 0.0032 ± 0.00012 min$^{-1}$, $K_{m,wt}$ = 3.3 ± 0.29 μM, $k_{cat,G226E}$ = 0.010 ± 0.00087 min$^{-1}$, and $K_{m,G226E}$ = 15.8 ± 2.5 μM.

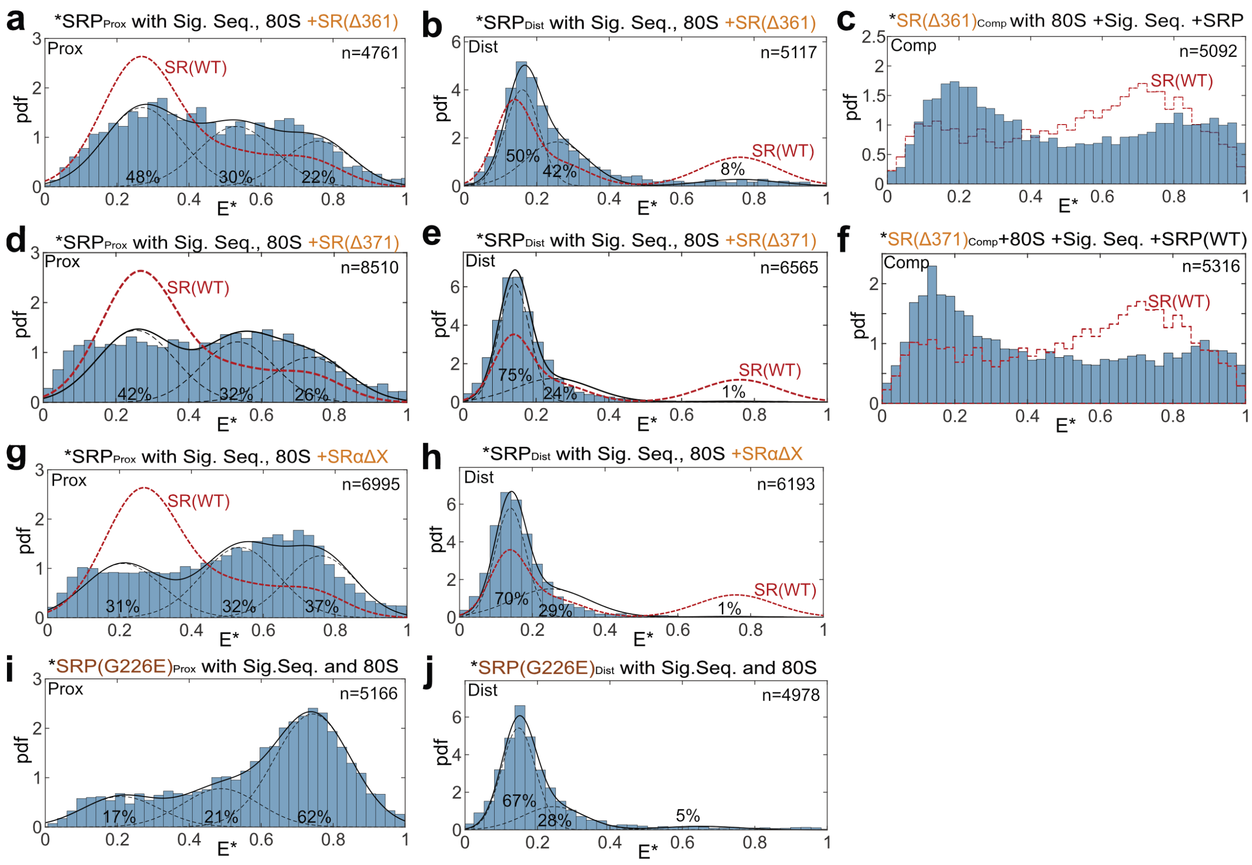


**Extended Data Figure 5**. The effect of SRP and SR mutations on the conformation of SRP and the SRP•SR complex. **a-f,** smFRET histograms of the targeting complex assembled with SR(Δ361) (a-c) or SR(Δ371) (d-f), detected by the Proximal (**a, d**), Distal (**b, e**), and SR compaction (**c, f**) probes. **g-h,** smFRET histograms of the targeting complex assembled with SRαΔX, detected using the Proximal (**g**) and Distal (**h**) probes. The red lines in (a)-(h) outline the data with the wildtype targeting complex. **i-j,** smFRET histograms of ribosome- and signal-sequence bound SRP(G226E), detected using the proximal (**i**) or distal (**j**) probes.

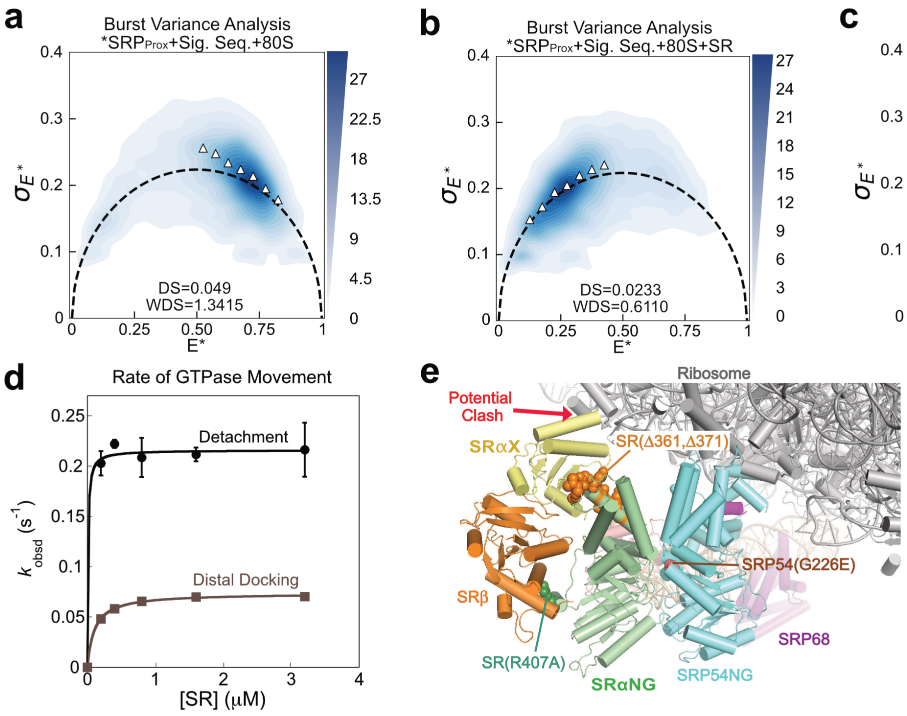


**Extended Data Figure 6.** Kinetic and structural analyses of the movement of the SRP54-NG•SR complex. **a-c,** Burst variance analysis of signal sequence- and ribosome-bound $SRP_{Prox}$ (a), $SRP_{Prox}$ in the targeting complex (b), and $SRP_{Dist}$ in the targeting complex (c). The black dashed curves represent the static limits. Triangles denote the SDs for individual E* bins ($SD_{E^*}$). **d,** The rate constants for movement of the SRP54-NG•SR complex from the ribosome exit site ('Detachment'), measured with SRP labeled with proximal probes, and for docking of this complex at the distal site ('Distal Docking') measured with the Distal probes. Changes in donor fluorescence intensity upon SR addition was monitored in real time on a stopped-flow apparatus to obtain the observed rate constants ($k_{obsd}$). Reactions contained 15 nM SRP-4A10L labeled with Proximal or Distal probes, 40 nM 80S, 200 μM GMPPNP, and the indicated SR concentrations. At saturating SR concentrations where $k_{obsd}$ is SR concentration-independent, $k_{obsd}$ is rate-limited by the unimolecular conformational change within the SRP•SR complex. Data are represented as mean ± SD, with n = 3-5. **e,** Structural model of the compacted SRP54-NG•SR complex at the ribosome exit site, generated by overlay of the SRP54-NG•SR complex in the distal state structure (PDB: 6FRK)[5] to SRP54-NG in the RNC•SRP complex (PDB: 3JAJ)[6]. The red arrow indicates potential clash with the ribosome. The locations of SRP and SR mutations are indicated.

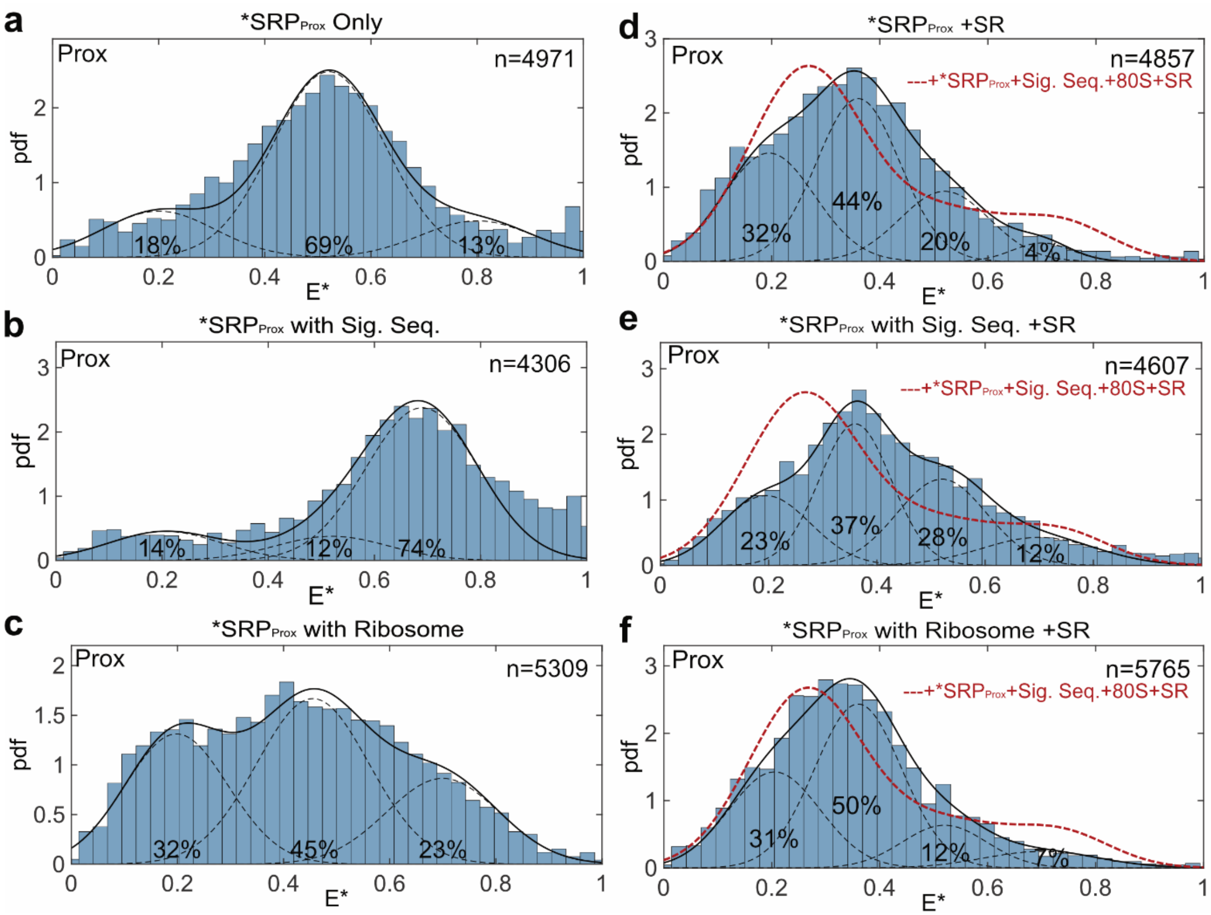


**Extended Data Figure 7**. Regulation of the conformational changes in SRP and the SRP•SR complex detected by the Proximal probes. **a-c,** smFRET histograms of $SRP_{Prox}$ in the absence (**a**) and presence of signal sequence (**b**) or ribosome (**c**). **d-f,** smFRET histograms of the $SRP_{Prox}$•SR complex in the absence (**d**) and presence of signal sequence (**e**) or ribosome (**f**). Reactions contained 100 pM $SRP_{Prox}$, 200 μM GMPPNP, 400 nM 80S, and 7 μM (**d**) or 1μM (**e**, **f**) SR where indicated. Reactions were incubated for 30-60 minutes before measurements to ensure that SRP-SR complex has formed. Red dashed lines outline the data for the complete targeting complex.

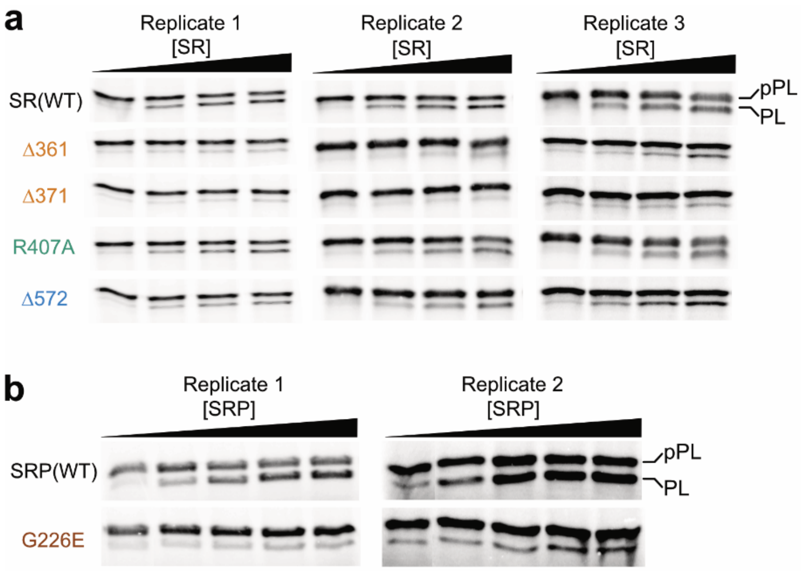


**Extended Data Figure 8**. SDS-PAGE autoradiographs for the targeting reactions mediated by the mutant SRP and SRs. Quantification of the translocation data are shown in Figures 4d and 4e.
