## Supplementary Methods for "Receptor compaction and GTPase movements drive cotranslational protein translocation"

***Preprotein targeting and translocation assays.***

Co-translational targeting and translocation of $^{35}$S-methionine labeled pPL into salt-washed, trypsinized rough ER microsome (TKRM)[1,2] were carried out as outlined in Extended Data Figure 3. Unless otherwise specified, translocation reactions contained 20 nM SRP (when [SR] is varied), 100 nM SR (when [SRP] is varied), and 0.5 eq of TKRM (as defined in [3]). The reactions in Extended Data Fig. 3 contained 7-Me-GTP, and 7-Me-GTP was omitted for the reactions shown in Figure 4 and Extended Data Fig. 8. Translocation efficiency was quantified as:

$$\text{Fraction Translocation} = \frac{(8/7)\text{prolactin}}{(8/7)\text{prolactin} + \text{preprolactin}}$$

The (8/7) term corrects for the difference in the number of methionines in preprolactin and signal-sequence cleaved prolactin.

***Measurement of GTPase rate constants.***

Basal GTPase reactions were measured under single-turnover conditions using varying concentrations of SRP54 in excess of γ-$^{32}$P-GTP. The SRP54 concentration dependence of the observed rate constant ($k_{\text{obsd}}$) were fit to Eq. 1,

$$k_{obsd} = k_{cat} \times \frac{[SRP]}{K_m + [SRP]} \qquad (1)$$

in which $k_{\text{cat}}$ is the basal GTPase rate constant at saturating SRP concentrations, and $K_{\text{m}}$ is the GTP concentration required to reach half of $k_{\text{cat}}$.

The reciprocally stimulated GTPase reaction between SRP and SR were measured under multiple turnover conditions using 0.2 μM SRP, varying concentrations of SR in excess of SRP, and 100 μM GTP (or 500 μM GTP for mutant SRP(G226E)) doped with trace γ-$^{32}$P-GTP. All reactions were measured in the targeting complex in which SRP was bound to the signal sequence and ribosome. The SR concentration dependences of the observed rate constants were fit to Eq. 2.

$$k_{obsd} = k_{cat} \times \frac{[SR]}{K_m + [SR]} \qquad (2)$$

As shown in Lee et al[2], the value of $k_{cat}/K_m$ is rate-limited by and therefore reports on the SRP-SR complex assembly rate constant.

***Steady state fluorescence measurements.***

Steady state fluorescence was measured using a Fluorolog 3-22 Spectrofluorometer (Horiba Jobin Yvon) following manufacturer's guidelines. To monitor the effects of SR, fluorescence emission spectra were recorded for 15 nM SRP-4A10L singly labeled with the indicated fluorescent dyes at specified positions, in the presence of 100 nM 80S with and without 1.5 μM SR present (Extended Data Fig. 2g-j). To monitor the effects of SRP, fluorescence emission spectra were recorded for 30 nM SR singly-labeled labeled with the indicated fluorescent dyes at specified positions, in the presence of 400 nM 80S with and without 400 nM SRP-4A10L present (Extended Data Fig. 2k-l). To monitor the effects of 80S, fluorescence emission spectra were recorded for singly labeled SRP-4A10L or SR in the absence and presence of 100 nM (for Proximal and Distal probes) or 400 nM (for Compaction probes) 80S (Extended Data Fig. 2m-p). All reactions also contained 200 μM GppNHp and 0.6 mg/ml BSA.

Fluorescence anisotropy was measured using the anisotropy module on Fluorolog 3-22 following the manufacturer's guidelines. The reaction contained 15 nM SRP-4A10L or 30 nM SR labeled with the indicated fluorescent dyes at specified positions (Extended Data Fig. 2q and r), 200 μM GppNHp, 0.6 mg/ml BSA, and 100 nM/400 nM 80S (for Distal/Compaction probes, respectively) where indicated.

Equilibrium titrations to measure the equilibrium dissociation constant ($K_d$) of the SRP•SR complex were carried out using 15 nM Cy3B labeled SRP-4A10L or SRP(G226E)-4A10L, 40 nM 80S, 2 mM GTP, and Atto647N-labeled SR at the indicated concentrations. SRP54 was labeled with Cy3B at C47, and SR was labeled at the C-terminus with Atto647N[2]. SR also contained the R458A mutation to block GTP hydrolysis and hydrolysis-driven dissociation of the SRP•SR complex[2]. Donor fluorescence intensity was recorded when equilibrium was reached. 0.6 mg/ml BSA was supplemented in SRP Assay buffer to reduce non-specific adhesion of proteins to surfaces. A control titration with unlabeled SR was carried out in parallel, and the fluorescence intensity change from the control reaction was subtracted. The fluorescence signal was converted to FRET (E) using Eq. 3,

$$E = 1 - \frac{F_{DA}}{F_D} \qquad (3)$$

where $F_{DA}$ and $F_D$ are the fluorescence signals with and without the acceptor present. Values of E were plotted against SR concentration and fit to Eq. 4

$$\Delta F = \frac{[SRP]+[SR]+K_d-[\sqrt{([SRP]+[SR]+K_d)^2-4\times[SRP]\times[SR]}}{2\times[SRP]} \qquad (4)$$

in which ΔF is the E normalized fluorescence, and $K_d$ is the equilibrium dissociation constant of the SRP•SR complex.

Rate constants of conformational changes in the targeting complex were measured using a stopped-flow apparatus (Kintek) by rapidly mixing 15 nM SRP-4A10L labeled with Proximal (Detachment) or Distal (Distal docking) probes with varying concentrations of unlabeled SR. The reactions also contained 40 nM 80S, 200 μM GppNHp, and 0.6 mg/ml BSA. The time courses of donor fluorescence intensity change were fit to exponential functions to extract the observed rate constants ($k_{obsd}$). The SR concentration dependences of $k_{obsd}$ were fit to Eq. 2 (Extended Data Fig. 6d) to extract the rate constant at saturation, which is not limited by SRP-SR complex assembly and reports on the rate of conformational change.

***smFRET measurements.***

Labeled SRP or SRP-4A10L was diluted to 100-200 pM in SRP Assay Buffer. Unless otherwise specified, the samples contained 200 μM GppNHp, 150 nM 80S, and 1.5 μM SRαβΔTM where indicated. To measure conformational changes in SR, SR labeled with compaction probes was diluted to 100-200 pM in SRP Assay Buffer containing 200 μM GppNHp, 400 nM 80S, and 400 nM SRP or SRP-4A10L where indicated. With signal sequence and ribosome-bound SRP, these concentrations are sufficient to ensure that all labeled species are bound with the indicated binding partners[2] (Fig. 4c). The reactions in Figure 3h and Extended data Fig. 7d contained 7 μM SR, and the reactions in Figures 3i,j and Extended data Fig. 7e,f contained 1 μM SR. The samples in these reactions were incubated at 25 °C for 30-60 minutes before measurements to ensure complete formation of the SRP•SR complex. Samples were placed in a closed chamber made by sandwiching a perforated silicone sheet (Grace Bio-Labs) with two coverslips to prevent potential evaporation during measurements. Data were collected over 30-60 min using an ALEX-FAMS setup with two single-photon Avalanche photodiodes (Perkin Elmer) and 532 nm (CNI laser) and 635 nm (Opto Engine LLC) continuous wave lasers operating at 150 μW and 70 μW, respectively[4,5].

All smFRET data analyses including burst search and burst selection were performed using FRETBursts[6], an open-source burst analysis toolkit for confocal smFRET. A dual-channel burst search[7] was performed to separate the particles containing FRET pairs from particles labeled with only the donor or acceptor dye. Each burst (assumed to be the fluorescence signal from an individual SRP or SR particle) was identified as a minimum of 10 consecutive detected photons with a photon count rate at between 6 to 15 times higher than the background photon count rate during both donor and acceptor excitation periods. Since the background rate can fluctuate within a measurement, the background rate was computed for every 50 second interval according to maximum likelihood fitting of the inter-photon delay distribution. The identified bursts were further selected according to the following criteria: (i) $n_{DD} + n_{DA} \geq 15$; and (ii) $n_{AA} \geq 15$, where $n_{DD}$ and $n_{DA}$ are the number of photons emitted from donor and acceptor during the donor excitation period, respectively, and $n_{AA}$ is the number of photons emitted from acceptor during the acceptor excitation period.

Under certain conditions, we observed some aggregated species that led to additional populations with different stoichiometry (S) values. The aggregated species are expected to show exceptionally long burst durations and/or multiple bursts that are close together if the dye molecules undergo photo-physical events such as blinking. To filter out such aggregates, we first fused bursts that are less than 5 milliseconds apart, then removed bursts that have burst durations larger than 10 milliseconds. After applying this filter, the FRET populations showed a single Gaussian distribution in terms of S-value. In rare cases, aggregated species dominated the FRET population, and those data were discarded and not included in any of the analyses.

The uncorrected FRET efficiency (E*) and Stoichiometry (S) for each burst were calculated using the following equations:

$$E^* = \frac{n_{DA}}{n_{DD} + n_{DA}} \qquad (5)$$

$$S = \frac{n_{DD} + n_{DA}}{n_{DD} + n_{DA} + n_{AA}} \qquad (6)$$

FRET histograms were obtained by 1D projection of the 2D E*-S histograms onto the E* axis. In most cases, E* is different from actual FRET efficiency due to simplifying assumptions (*i.e. lk=0*, *dir=0*, *γ=1).* However, since the correction factors only depend on the photo-physical properties of the fluorophores and the configuration of the optical setup, their contributions to FRET efficiency are constant as long as the same optical set up, FRET pair and labeling position are used throughout all measurements. Importantly, we did not observe any significant changes in the photo-physical properties of both the donor and acceptor dyes due to local environments (i.e. different conformations, substrates, ligands) (Extended Data Fig. 2). Therefore, conformational changes in SRP that alter the actual FRET efficiency will also change the E* value, and the trend of the changes with different binding partners will be the same.

*Burst Variance Analysis.*

Burst Variance Analysis (BVA)[8] was implemented to investigate conformational dynamics of SRP. Variance in fluorescence signal larger than the shot-noise limit can be attributed to static heterogeneity (i.e. mixture of stable multispecies with similar FRET values), dynamic heterogeneity (i.e. one species dynamically transitioning between states), or combination of dynamic and static heterogeneity. BVA can identify the dynamics of molecules by comparing the empirical standard deviation (SD) of E* of sub-bursts (containing fixed number of photons, *n*) within a burst to the expected shot-noise limited SD (i.e. Static limit). Eq. 7 was used to compute the static limit for a given mean E*.

$$\text{Static limit } (n, E^*) = \sqrt{\frac{E^*(1-E^*)}{n}} \qquad (7)$$

where $n = n_{DD} + n_{DA}$, the number of photons in a sub-burst. In this study, we used $n$ = 5. The static limits were represented as dashed curves in BVA plots (Fig 2n, Extended Data Figure 6a-c**)**. The observed SD ($\sigma_{E^*}$) for individual molecules was computed using Eq. 8.

$$\sigma_{E^*} = \sqrt{\frac{1}{M_i}\sum_{j=1}^{M_i}(e_{ij}^* - E_i^*)^2} \qquad (8)$$

where $M_i$ is the number of sub-bursts in the $i$ th burst, $e_{ij}^*$ is an uncorrected FRET efficiency of the $j$th sub-burst, and $E_i^*$ is the mean of all sub-bursts' uncorrected FRET efficiencies in the $i$ th burst. To reduce error that could arise from individual bursts, which only contain a small number of sub-bursts, we first binned bursts along the $E^*$ axis into 20 bins with a bin width of 0.05. All of the sub-bursts in each bin within bursts were then used to calculate the observed SD for each bin ($SD_{E^*}$) using Eq. 9,

$$SD_{E^*} = \sqrt{\sum_{\substack{i \text{ where} \\ L \le E_i^* < U}} \sum_{j=1}^{M_i} \left[ \frac{(e_{ij}^* - \mu)^2}{\sum M_i} \right]} \qquad (9)$$

where $\mu = \sum_{\substack{i \text{ where} \\ L \le E_i^* < U}} \sum_{j=1}^{M_i} \left( e_{ij}^* \Big/ \sum M_i \right)$, $L$ is the lower bound of the bin, $\mu$ is the upper bound of the bin, $M_i$ is the number of sub-bursts in the $i$ th burst, and $e_{ij}^*$ is the uncorrected FRET efficiency of the $j$ th sub-burst in the $i$ th burst**.**

The dynamic score (DS) was computed using Eq. 10.

$$DS = \sqrt{\sum_{SD_{E^*} - SD_{E^*,\text{static}} > 0} (SD_{E^*} - SD_{E^*,\text{static}})^2} \qquad (10)$$

where $SD_{E^*,\text{static}}$ is the $SD_{E^*}$ of simulated static molecules using the Monte Carlo method and $N_{E^*}$ is the number of bursts in a given bin. The weighted dynamic score (WDS) was computed using Eq. 11 to take into account the different numbers of bursts in each bin.

$$WDS = \sqrt{\sum_{SD_{E^*} - SD_{E^*,\mathrm{static}} > 0} \left( \frac{N_{E^*}}{\sum N_{E^*}} \right) (SD_{E^*} - SD_{E^*,\mathrm{static}})^2} \qquad (11)$$

To ensure that the score reports on the dynamics of the major species in a given sample, only the $SD_{E^*}$ for bins with at least 5% of the total number of bursts in the entire FRET histogram (denoted by triangles in BVA plots in Fig 2n and Extended Data Figure 6a-c**)** were used to calculate DS and WDS.
